# Adaptation shapes local cortical reactivity: from bifurcation diagram and simulations to human physiological and pathological responses

**DOI:** 10.1101/2022.06.11.493219

**Authors:** Anna Cattani, Andrea Galluzzi, Matteo Fecchio, Andrea Pigorini, Maurizio Mattia, Marcello Massimini

## Abstract

Human studies employing intracerebral and transcranial perturbations suggest that the input-output properties of cortical circuits are dramatically affected during sleep in healthy subjects as well as in awake patients with multifocal and focal brain injury. In all these conditions, cortical circuits react to direct stimulation with an initial activation followed by suppression of activity (Off-period) that disrupts the build-up of sustained causal interactions typically observed in healthy wakefulness. The transition to this stereotypical response is of clinical relevance, being associated with loss of consciousness or loss of function. Here, we provide a mechanistic explanation of these findings by means of mean-field theory and simulations of a cortical-like module endowed with activity-dependent adaptation. First, we show that fundamental aspects of the local responses elicited in humans by direct cortical stimulation can be replicated by systematically varying the relationships between adaptation strength and excitation level in the network. Then, we reveal a region in the adaptation-excitation parameter space of key relevance for both physiological and pathological conditions, where spontaneous activity and responses to perturbation diverge in their ability to reveal Off-periods. Finally, we substantiate through simulations of connected cortical-like modules the role of adaptation mechanisms in preventing cortical neurons from engaging in reciprocal causal interactions, as suggested by empirical studies. These modeling results provide a general theoretical framework and a mechanistic interpretation for a body of neurophysiological measurements that bears key relevance for physiological states as well as for the assessment and rehabilitation of brain-injured patients.

**Significance Statement:** Suppression of cortical activity following an initial activation is a defining feature of deep sleep in healthy subjects and wakefulness in patients affected by focal and multifocal brain injuries. Experimental findings suggest that these bimodal responses disrupt the emergence of complex interactions among cortical regions, leading to loss of consciousness or functional impairments. Given their practical implications, it is important to study the mechanisms involved within a general theoretical framework. Using a neuronal network model, we provide evidence for a key role of activity-dependent adaptation mechanisms in shaping the responses to perturbation and in affecting the build-up of complex cortical interactions. Overall, this work provides a mechanistic interpretation relevant for the stratification, follow-up, and rehabilitation of brain-injured patients.

## Introduction

Activity-dependent adaptation accounts for local fatigue mechanisms, i.e., self-inhibition, which lowers the firing rate of neuronal populations. Adaptation can be induced by several microscopic biophysical mechanisms mainly involving muscarinic and calcium-dependent potassium channels (Gerstner et al., 2014). As suggested by mean-field theories, activity-dependent adaptation is key for the generation of the rhythmic alternation between On- (high-activity Up state) and Off- (low-activity Down state) periods occurring during slow-wave sleep in excitable cortical networks (Latham et al., 2000; Gigante et al., 2007). Indeed, this fatigue mechanism accumulates during the On-period, destabilizes self-sustained synaptic reverberation, and eventually drives the neuronal population toward the Off-period. During the Off-period, due to the low neuronal firing, fatigue relaxes and the network can transition to the ensuing active On-period (Compte et al., 2003; Mattia and Sanchez-Vives, 2012).

In human recordings, the spontaneous occurrence of Off-periods can be inferred through the detection of slow waves associated with brief suppressions of high frequency power (Mukovski et al., 2007), as shown in scalp (Piantoni et al., 2013) and intracranial measurements of ongoing activity during sleep and anesthesia (Cash et al., 2009; Lewis et al., 2012). In the context of empirical neuroscience, the tendency of cortical circuits to fall into an Off-period is often referred to with the term *cortical bistability* (Tononi and Massimini, 2008; Nir et al., 2011). This term does not necessarily coincide with the notion of bistability in nonlinear physics and dynamical systems theory, but it is rather used to indicate the fact that circuits are prone to plunge into a silent Off-period following an active On-period.

Recent studies employing direct cortical perturbations, such as transcranial magnetic stimulation (TMS) (Massimini et al., 2007) and intracranial electrical stimulations (Pigorini et al., 2015; Usami et al., 2015), have demonstrated that cortical bistability is a common feature characterizing cortical responsiveness during non-rapid eye movement sleep (NREM) and in different pathological conditions. Indeed, cortical circuits react to perturbations with an Off-period also in vegetative state patients with severe brain injuries (Rosanova et al., 2018), as well as in the perilesional area surrounding focal cortical lesions in stroke patients (Sarasso et al., 2020; Tscherpel et al., 2020). Remarkably, in all these conditions perturbations can elicit clear-cut Off-periods and reveal bistability even when this is not evident in the spontaneous EEG. Further, the same studies show that the occurrence of Off-periods can disrupt the emergence of complex spatiotemporal patterns, bearing profound clinical implications. When cortical bistability involves most of the cortex, such as in the unresponsive wakefulness syndrome, large-scale brain interaction collapse leading to loss of consciousness (Rosanova et al., 2018), whereas local Off-periods, such as those found in the perilesional areas of stroke patients, are associated with regional circuit impairment and selective functional deficits (Sarasso et al., 2020).

Given the practical relevance of these findings, it is important to understand cortical bistability as revealed by responses to direct cortical perturbations, its mechanisms, its potential dissociation from spontaneous activity and its impact on cortico-cortical interactions within a general formal framework. In this work, we do so by adopting a theoretical approach that entails a cortical module endowed with activity-dependent adaptation (Mattia and Sanchez-Vives, 2012). Here, we first perform a bifurcation analysis of the model to classify its dynamical regimes. Then, we explore how the spontaneous and stimulus-evoked dynamics of the cortical module vary parametrically as a function of adaptation and excitation levels. Further, within this theoretical framework, we turn to explain the empirical finding that direct cortical stimulation is more effective than the observation of ongoing dynamics in revealing Off-periods in both physiological and pathological conditions. Finally, we consider two linked modules and we study how changing the adaptation level can affect their ability to engage in reciprocal interactions without resorting to any change in their connectivity.

## Materials and Methods

### Spiking neuron network

The neuronal network model is adapted from (Torao-Angosto et al., 2021) where it has been proven to quantitatively reproduce the statistical features of Up-Down slow oscillations recorded in L5 of the visual cortex of sleeping and anesthetized rats. Briefly, the network is composed of 6300 excitatory (E) and 2580 inhibitory leaky integrate-and-fire (LIF) neurons. The membrane potential *V*_*i*_ of the *i*-th neuron evolves as

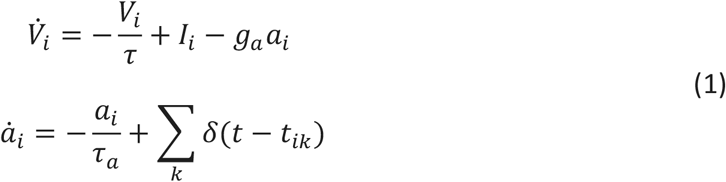

emitting its *k*-th spike at time *t*_*ik*_ if *V*_*i*_(*t*_*ik*_) ≥ *V*_*thr*_ for the first time from the emission of its previous action potential. Following the spike emission, *V*_*i*_ = *V*_*res*_ for a refractory absolute period *τ*_0_ [2 ms (1 ms) for E (I) neurons] before restarting its time evolution (1). Reset potential *V*_*res*_ = 15 mV and the emission threshold *V*_*thr*_ = 20 mV are the same for both neuron types, while decay constant *τ* is 20 and 10 ms for E and I neurons, respectively. The synaptic current 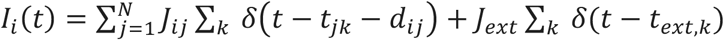 results from presynaptic spiking activity mediated by recurrent synaptic efficacies *J*_*ij*_ randomly selected to be different from zero with connection probability *c*_*EE*_, *c*_*EI*_, *c*_*IE*_, *c*_*II*_ = {0.6, 5, 0.2, 1.7}%. *J*_*ij*_ takes positive or negative values according to the type of of the *j*-th presynaptic neuron, and it is randomly extracted from a truncated Gaussian distribution with mean *J*_*EE*_, *J*_*EI*_, *J*_*IE*_, *J*_*II*_ = {1.9, −1.1, 2.2, −1.1} mV and a relative standard deviation of 25%. Presynaptic spikes are delivered with an axonal delay *d*_*ij*_ randomly sampled from an exponential distribution with average of 22.6 ms and 5.7 ms for E and I presynaptic neurons, respectively, modeling non-instantaneous synaptic transmission (Mattia et al., 2019). External neurons contribute to *I*_*i*_(*t*) as a Poissonian spike train made of *C*_*ext*_ independent sources each with an average firing rate of 0.25 Hz. Here, for E neurons *C*_*ext*_ range from 3200 and 3350 in the bifurcation diagram of Fig. 1, while *C*_*ext*_ = 733 for I neurons. External spikes occurring *t*_*ext,k*_ affect *I*_*i*_(*t*) with efficacies *J*_*ext*_ = 0.48 mV and 2.2 mV for E and I neurons, respectively.

**Figure 1.**
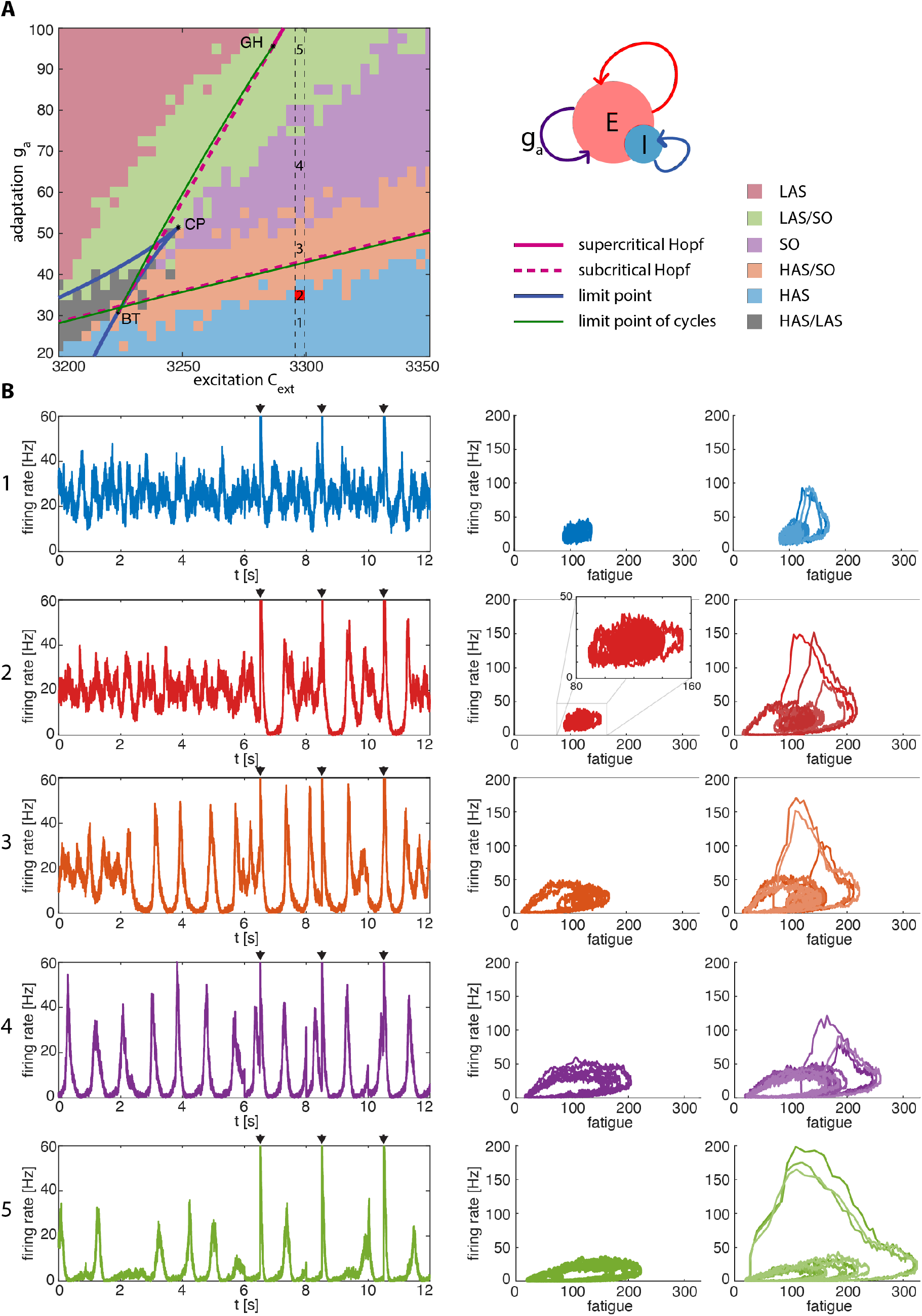
Bifurcation analysis of the population rate model and dynamical regimes of the spiking neuron network. **A:** codimension-two bifurcation diagram in the excitation-adaptation parameter space, namely (*C*_*E,ext*_, *g*_*a*_), superimposed to the dynamical regimes of the spiking neuron network classified according to its spontaneous activity (see section “Analysis of the dynamical regimes of the spiking neuron network” in Materials and Methods for details). The codimension-one bifurcations encompass the subcritical Andronov-Hopf bifurcation curves (dashed magenta lines), the supercritical Andronov-Hopf bifurcation curves (solid magenta line), the saddle-node bifurcation curve (blue curves), and the limit point of cycles (green lines) that turn the limit cycles originating from Andronov-Hopf stable. The codimension-two bifurcation Bogdanov-Takens (BT) is the contact point between the saddle-node bifurcation curve and Andronov-Hopf bifurcation curves. The saddle-node curve shows a cusp bifurcation (CP). The areas among the curves have been color-coded according to the regime shown by the spontaneous dynamics of the spiking neuron network. The five different dynamical regimes are characterized by HAS (blue area), HAS with incursions of Off-periods due to finite-size effects (orange area), SO (purple area), LAS with incursions of On-periods (green area), and, finally, LAS (green areas). A sixth region (grey area), not of interest in the present work, is characterized by HAS and LAS alternating at an irregular pace. **B:** Spontaneous and stimulus-evoked signals using the same color-coding as in A. Left column, time series encompassing both spontaneous activity (up to 6000 time steps) and stimulus-evoked activity (remaining interval). Black triangles indicate the occurrence of the stimulation. Central column, spontaneous activity as a function of fatigue. Right column, three superimposed orbits due to perturbations as a function of the fatigue.

E-neurons incorporate the activity-dependent fatigue mechanism modeling spike-frequency adaptation (SFA). To this aim, the dynamics of the adaptation level *a*_*i*_(*t*) modulates a hyperpolarizing current (Gigante et al., 2007; Mattia and Sanchez-Vives, 2012). The higher the adaptation level (calcium concentration) due to the spikes emitted by the *i*-th neuron, the higher is its self-inhibition eventually reducing its firing rate. This fatigue mechanism mimics the extracellular ionic concentrations which drive a hyperpolarizing potassium current, modulated by the adaptation strength *g*_*a*_ ranging from 20 to 100 mV/s. Relaxation time constant for the adaptation level is *τ*_*a*_ = 150 ms.

We also performed simulations of a network made by two identical modules. Each module is made by E- and I-neurons as described above, with the only exception of a zero relative standard deviation for synaptic efficacies. The two modules interact with each other through connections established between excitatory neurons only, with synaptic efficacy set to 1.18 mV. The probability of connection for each neuron is 0.1%, and average axonal delays is set to 55 ms for connections from the first to the second module, and to 50 ms from the second to the first module.

Simulations of the spiking neuron network were performed with an open source program relying on an event-driven numerical integration described in (Mattia and Giudice, 2000) and available as open source software from (https://github.com/mauriziomattia/Perseus). A detailed explanation of the spiking neuron network used in the present work can be found on the EBRAINS Collaboratory (*request submitted to EBRAINS, DOI currently pending*) in the form of a Jupyter Notebook.

### Population rate model

Under mean-field approximations requiring many presynaptic contacts *c*_*αβ*_*N* and limited firing rates *v*_*α*_ (Amit and Tsodyks, 1991; Amit and Brunel, 1997), the evolution of the instantaneous firing rate *v*_*α*_(*t*) of an infinite-size networks is described by 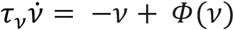 (Treves, 1993; Brunel and Hakim, 1999; Mattia and Del Giudice, 2002) (Treves, 1993; Brunel and Hakim, 1999; Mattia and Giudice, 2000). Here the current-to-rate gain function *Φ* is the Siegert-Ricciardi one (Siegert, 1951; Capocelli and Ricciardi, 1971):

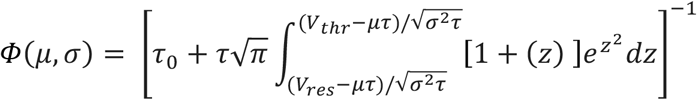

where *μ* and *σ*^2^ are the infinitesimal mean and variance of the input current, respectively, and *erf*(x) is the error function. Since the cortical module we are considering includes two interacting populations of excitatory and inhibitory neurons, the mean-field dynamics is

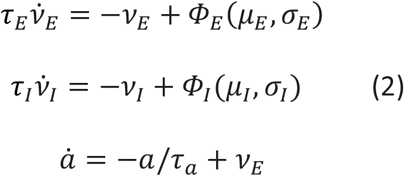

where *α* ∈ {*E, I*}, *Φ*_*α*_(*μ*_*α*_, *σ*_*α*_) is the above gain function computed with the single-neuron parameters of type *α* and *τ*_*α*_ is the decay constant of the LIF membrane potential. The infinitesimal mean and variance of the input current are defined as:

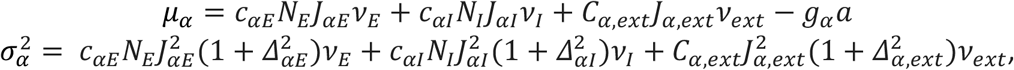

with *α, β* ∈ {*E, I*} and *Δ*_*αβ*_ are the standard deviations of the synaptic efficacies *J*_*ij*_ for the different neuronal types. This mean-field approximation provides an effective description of the collective dynamics of networks of LIF neurons with spike-frequency adaptation even far from equilibrium (Amit and Tsodyks, 1991; Amit and Brunel, 1997). Here only excitatory neurons have adaptation: *g*_*E*_ ≡ *g* and *g*_*I*_ = 0.

Finally, to compute the bifurcation diagram in Fig. 1, we further reduced the dimensionality of the rate equation (2) to 2 by assuming the limit of a quasi-adiabatic approximation (*τ*_*I*_ → 0) as in (Mascaro and Amit, 1999). Due to the breaking of the diffusion approximation valid only in the limit *J*_*αβ*_ → 0, we shifted horizontally the critical point of this bifurcation diagram by a suited amount (*ΔC*_*E,ext*_ = −*310*) of excitation level. Indeed, in the diffusion limit the currents received by the neurons are continuous stochastic processes determined by the continuous barrage of the synaptic input arriving at high rates and inducing small jumps in the membrane potential (Tuckwell, 1988). However, in the model network we simulated, synaptic efficacies are not negligible and the membrane potentials have a jump-like evolution in time. In this shot-noise regime, firing rates are smaller compared to the ones predicted under diffusion approximation (Richardson and Swarbrick, 2010), and this explains why we need to incorporate this effect by reducing the excitation level in our mean-field theory eventually leading to a rightward shift of the theoretical bifurcation diagram. Furthermore, we also shifted vertically the critical points of the bifurcation diagram by a suited amount (*Δg*_*a*_ = −*7* mV/s) of adaptation. Besides the breaking of the diffusion approximation, the shifts *ΔC*_*E,ext*_ and *Δg*_*a*_ are expected to be theoretically justified by the fact that the simulated networks are composed of a finite number of spiking neurons challenging the infinite-size limit upon which extended mean-field approximation relies on (Mattia and Del Giudice, 2002).

### Simulated data and data preprocessing

Simulations consist of N trials (N = 50 for the single cortical module, and N = 250 for the case of two interacting modules). The external stimulus in each trial was delivered at time t=0, with a pre-stimulus interval lasting randomly between 5000 and 5300 ms, and post-stimulus interval lasting 5000 ms. Stimulation artifact was reduced by applying a Tukey-windowed median filtering, as in (Chang et al., 2012), between -5 and 5 ms (Pigorini et al., 2015).

### Bifurcation analysis of the population rate model

A detailed analysis of local bifurcations of Eq. 2 (reduced to 2 equations under the assumption of a quasi-adiabatic approximation (*τ*_*I*_ → 0)) was performed through MatCont (Dhooge et al., 2003), a Matlab Toolbox that allows computing curves of equilibria and bifurcation points through a prediction-correction continuation algorithm, as described in (Kuznetsov, 2004).

### Analysis of the dynamical regimes of the spiking neuron network

The dynamical regimes of the spiking neuron network (Eq. 1) were determined according to the frequency of On and Off-periods (Up and Down states, respectively) detected across 50 trials of spontaneous activity lasting 2 seconds each. The detection of the On and Off-periods was carried out by looking at the firing rate in each trial. If the firing rate exceeded 20 Hz, then the trial contained at least one On-period. On the contrary, if the firing rate fell under 5 Hz, the trial contained at least one Off-period. Once we collected the probability of On and Off-periods across 50 trials for each value of excitation and adaptation level, we classified five dynamical regimes of interest. Following the nomenclature in (Gigante et al., 2007; Mattia and Sanchez-Vives, 2012), high-asynchronous state (HAS) is characterized by a probability of On-periods equal to 1 (i.e., sustained high firing rate in every trial) and absence of Off-periods. Low-asynchronous state (LAS) presents a probability of Off-periods equal to 1 (i.e., persistent low firing rates in every trial) and absent On-periods. Slow oscillations state (SO) is characterized by both probability of On and Off-periods equal to 1 since each trial sees On and Off-periods alternating at a regular pace. Furthermore, on the border between LAS and SO, there is a region where the probability of Off-periods is equal to 1 with incursions of On-periods states (non-zero probability of On-periods < 1). Vice versa, the region between HAS and SO is characterized by On-periods in all the trials with incursions of Off-periods (non-zero probability of Off-periods <1). Finally, a sixth region, which is not the focus of this work, encompasses both HAS and LAS with non-zero probability of On and Off-periods lower than 1. This is the case of heterogeneous trials, some containing only On-periods, some only Off-periods, and some exhibiting both On and Off-periods.

### Data analysis in the power and phase domain

Data analysis was performed using Matlab R2020b (The MathWorks Inc.). To analyze the firing rate responses in the time-frequency domain, we used the event-related spectral perturbation (ERSP) implemented in EEGLAB (Delorme and Makeig, 2004). More in detail, single trials were time-frequency decomposed between 5 Hz and 30 Hz using Wavelet transform (Morlet, window span: 3.5 cycles). The resulting ERSPs were averaged across trials, and eventually normalized by subtracting the mean baseline power spectrum (from −5000 ms to -400 ms). To detect statistically significant activation in the time–frequency domain, we applied a bootstrap statistics (α < 0.005), with a number of permutations set to 1000. Power values that were not significantly different from the baseline were set to zero.

To quantify the coherence of the response to the perturbation across trials, we quantified the inter-trial coherence (ITC, also referred as inter-trial phase-locking factor) (Delorme and Makeig, 2004) at all the frequencies in between 5 Hz and 30 Hz. Furthermore, we retained only the ITC values that correspond to time and frequency bins with significant increase of power, and we summed the non-zero ITC values over the frequencies. To the extent that the ITC measures the coherence of the response to a perturbation across trials in a specific time window, it can be used to quantify the duration of the deterministic effect of a given external input.

## Results

We employ a spiking network model to investigate to what extent varying the strength of adaptation and excitation can explain fundamental dynamics of cortical responsiveness empirically observed in healthy humans and brain-injured patients. Specifically, we aim at replicating and providing a mechanistic explanation of experimental data encompassing TMS-EEG recordings in healthy subjects during N2 sleep (Massimini et al., 2007), in VS/UWS patients with severe brain injury (Rosanova et al., 2018), in stroke patients (Sarasso et al., 2020; Tscherpel et al., 2020), as well as intracranially-evoked potentials in wakefulness/sleep (Pigorini et al., 2015).

### Bifurcation analysis and characterization of the dynamical regimes

Activity-dependent adaptation and excitation level shape the ongoing and stimulus-evoked activity of the simulated cortical module (Eq. 1). To fully characterize the dynamical regimes the model accounts for, we first performed a bifurcation analysis of its continuous counterpart (Eq. 2) exploiting a numerical continuation technique (Dhooge et al., 2003; Kuznetsov, 2004). This analysis expands previous studies (Gigante et al., 2007; Mattia and Sanchez-Vives, 2012) providing a full characterization of the bifurcations of the mean-field dynamics of the network. Taking into account only two key parameters, i.e., the level of excitation (*C*_*E,ext*_) and adaptation (*g*) (see Materials and Methods), the bifurcation diagram in Fig. 1A shows critical points such as *limit points* (blue traces) and *subcritical Hopf bifurcations* (dashed magenta traces). Subcritical Hopf bifurcations lie near the *limit point of cycles* (green traces) where stable limit cycles arise. Thus, in the region limited by these two curves, stable slow oscillations are generated, corresponding to the cortical bistability observed in neurophysiological recordings performed in anesthetized rats (Tort-Colet et al., 2021). On the contrary, the region enclosed by the limit point curve (blue trace) is characterized by two stable and one unstable equilibrium solutions. The coexisting stable equilibria are characterized by high and low asynchronous firing states (HAS and LAS, respectively) successfully modeling the cognitive function of working memory (Amit and Brunel, 1997). In order to disambiguate, it is worth noting that this regime, which is not the focus of the present work, is also indicated with the term “bistable” as classically understood in the field of dynamical systems (Gerstner et al., 2014).

To check the effectiveness of this bifurcation diagram, we characterized the spontaneous dynamical regimes of equivalent networks of spiking (adaptive leaky integrate-and-fire) neurons. The leftmost part of the traces in Fig. 1B (left column) shows 6 seconds of spontaneous activity for a given excitation value and five different adaptation levels. By taking into account the probability of Off and On-periods during spontaneous activity (see Materials and Methods for details), we identified different regions. The light blue region displays HAS, i.e., relatively high firing rate and absence of Off-periods. The light red region is characterized by LAS, i.e., low asynchronous firing rate distinctive of Off-periods and absence of On-periods. The purple region features a regular alternation between HAS and LAS at a regular pace generating slow oscillations (SO). The transition between these regimes is not as sudden as in the population rate model while varying one or both parameters. Indeed, finite-size effects make a state to be dominant, with incursions of the state of the nearby region (Gigante et al., 2007; Mattia and Sanchez-Vives, 2012). This is the case of the orange and green regions characterized by dominant HAS and LAS, respectively, with incursions of SO.

### Perturbations reveal cortical bistability beyond spontaneous activity

Crucially, we further characterized the dynamical regimes of the spiking neuron model (Eq. 1) by exploring its responses to perturbations, which consist of 1 ms-lasting external current injected into the excitatory neurons resembling the effects of direct magnetic and electrical stimulation. The role of the perturbation is to transiently alter the spontaneous population firing rate, potentially unveiling interesting activity-dependent properties of the system. The rightmost part of the traces in Fig. 1B (left column) shows 6 seconds of the stimulus-evoked activity for a given excitation value and five different adaptation levels. In the middle and right columns, the firing rate traces have been represented as a function of the average fatigue determining the adaptation of spike rates for both the spontaneous and the stimulus-evoked activity.

In case 1, the stimulus-evoked activity and spontaneous activity do not show relevant dissociations, as both are characterized by the absence of Off-periods, as empirically observed during wakefulness in human data. Higher adaptation levels, corresponding to cases 3-5, are all characterized by the presence of Off-periods in both the spontaneous and evoked activity, whose duration increases with the adaptation level, qualitatively resembling N3 sleep (case 3), anesthesia (case 4), and burst suppression (case 5) in human data. Crucially, case 2 in Fig. 1B reveals a clear-cut dissociation between the dynamical features observed in the spontaneous activity and in the stimulus-evoked response. Here, while Off-periods are not detectable in the spontaneous activity, they can be reliably triggered by perturbations. Hence, the model shows the existence of a particular regime whereby only an interventional approach with direct stimulations can reveal the underlying state of the system and its position in the bifurcation graph. Such dissociation between observable dynamics and responses to perturbations is important because it reproduces a key feature reported by empirical works. This is illustrated in Fig. 2 where the results of simulations are directly compared to the results of TMS-EEG experiments both in the time domain and in the frequency domain to detect the presence of cortical bistability (slow waves and Off-periods). This analysis shows a fundamental correspondence between case 2 of the model (Fig. 2A) and spontaneous as well as evoked patterns of electrophysiological activity found in humans across different conditions encompassing N2 sleep (Massimini et al., 2007) (Fig. 2B), UWS patients (Rosanova et al., 2018) (Fig. 2C), and perilesional area of stroke patients (Sarasso et al., 2020) (Fig. 2D). In all these cases, TMS-evoked slow waves and Off-periods, as detected by a significant high-frequency (>20 Hz) power suppression, are reliably revealed by cortical perturbations, whereas they are not visible in the spontaneous pre-stimulus activity. Considering its prevalence in real data across different physiological and pathological conditions, we asked whether such dissociation between spontaneous and stimulus-evoked activity is a peculiar feature of the specific level of excitation, or whether it can be generalized to a large interval of excitation levels. Fig. 3 shows the difference in the frequency of Off-periods detected by analyzing the spontaneous and evoked activity for each excitation-adaptation pair. We found an area surrounding the low Hopf-bifurcation curve, extending to about 10% of the analyzed excitation-adaptation plane, characterized by a pronounced difference between evoked and spontaneous Off-periods. More specifically, despite a marked difference being present for all the excitation levels higher than *C*_*E,ext*_ = 3250, a near-complete dissociation (difference close to 1) can be shown for very high excitation levels.

**Figure 2.**
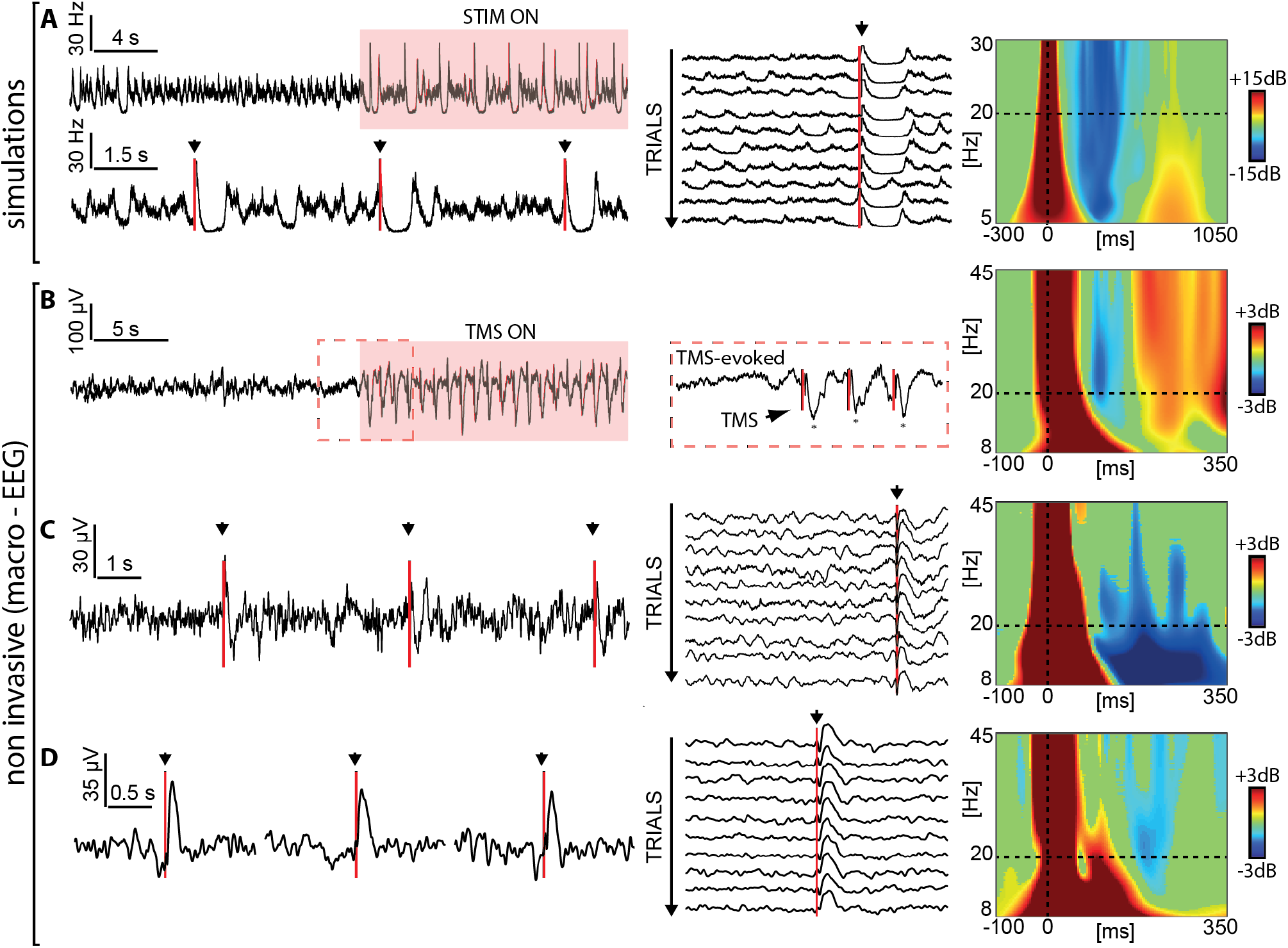
Perturbations trigger an Off-period regardless of spontaneous baseline activity in both simulated data, with adaptation level sets as in case 2 of Fig. 1, and in TMS-EEG data (during physiological N2 sleep, UWS patients, and in the perilesional area of stroke patients). First column: time series with a few stimulations. Second column: trials without Off-periods in the baseline aligned to the stimulation onset. Third column: Event-related spectral perturbation (ERSP). Blue color indicates a significant reduction compared to the baseline, while red indicates a significant increase. Black triangles and red lines indicate the occurrence of the stimulation. **A**: simulated firing rate data. **B**: re-edited from (Massimini et al., 2007). EEG signal recorded from a channel (Cz-re-referenced) located under the stimulator during one TMS-ON block over a background of spontaneous NREM sleep (single-subject data). The TMS-ON block consisted of 40 stimuli at 0.8 Hz. The red dashed sections show the slow waves triggered at the beginning and at the end of the block. **C**: re-edited from (Rosanova et al., 2018). EEG recordings during TMS stimulations in a UWS patient. EEG activity of one representative electrode -Cz-re-referenced while TMS was delivered with an inter-stimulus interval randomly jittering between 5000 and 5300 ms. Middle panel, trials aligned to the stimulation displaying baseline activity without spontaneous slow waves. Panel **D** is derived from published data presented in Table 1 and Fig. 2 in (Sarasso et al., 2020). EEG recordings during TMS stimulations in a middle cerebral artery ischemia patient (patient 4). EEG activity of a contact (17) in the perilesional area re-referenced to average reference while TMS was delivered with an inter-stimulus interval randomly jittering between 2000 and 2300 ms. ERSPs in B, C, and D were computed as in (Rosanova et al., 2018).

**Figure 3.**
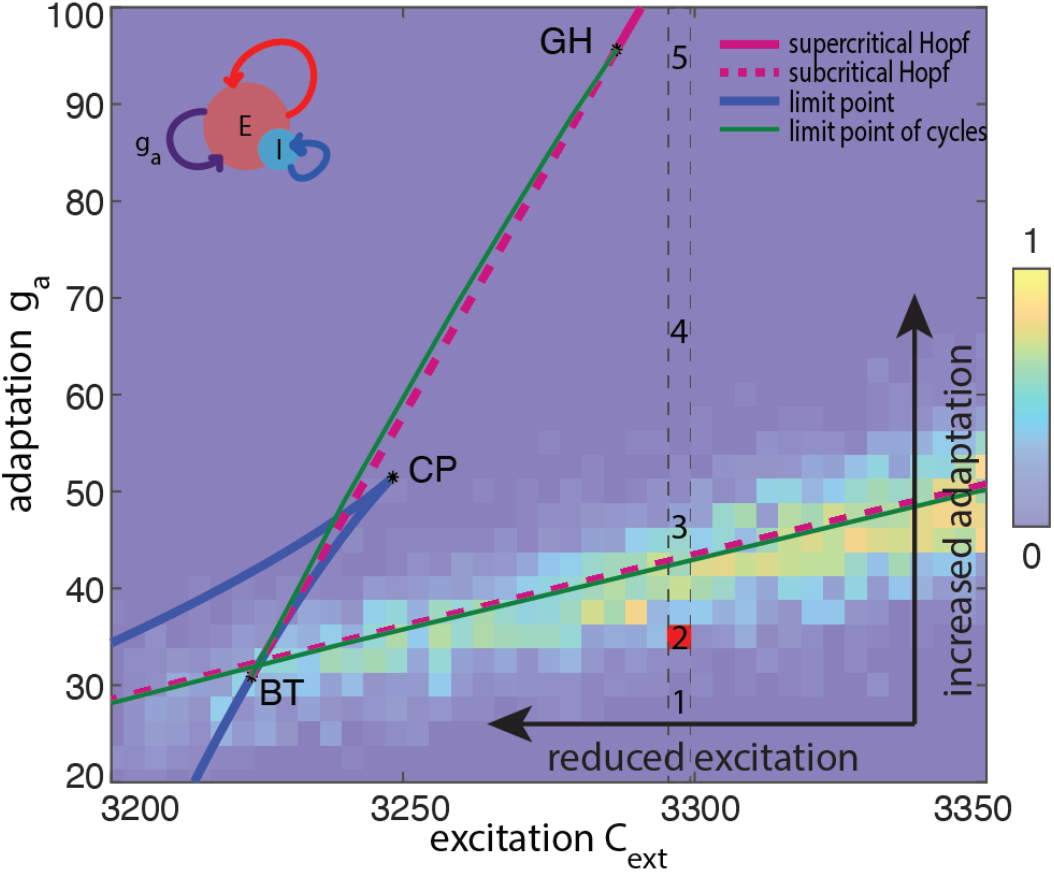
Difference between the fraction of evoked Off-periods and spontaneous Off-periods for each excitation-adaptation pair, superimposed on the bifurcation diagram of the population rate model shown in Fig. 1. The fraction of Off-periods has been computed over 100 trials lasting 2 seconds each.

Given the shallow slope of the lower branch of the Hopf bifurcation, changes in adaptation are expected to have more profound effects on cortical bistability than shifts in the excitation level. However, substantial changes of excitation levels for a given adaptation (i.e., horizontal shifts towards the left in the bifurcation diagram) still result in changes of spontaneous and stimulus-evoked dynamics similar to those reported in Fig. 1. Fig. 1-1 illustrates such cases, including the interesting instance (case 2) characterized by an intermediate excitation level, showing the dissociation between spontaneous and stimulus-evoked dynamics.

### High adaptation level is associated with early breakdown of the causal interactions compared to low adaptation

Empirical studies employing perturbations in humans (Pigorini et al., 2015; Usami et al., 2015; Rosanova et al., 2018), rodents (Arena et al., 2021) and cortical slices (D’Andola et al., 2018) suggest that cortical bistability and the associated Off-periods can disrupt the emergence of sustained patterns of causal interactions among cortical neurons. We thus used the model to test whether changes in adaptation level can also affect casual interaction among groups of cortical neurons. To do that, we considered two cortical modules, instead of one as in previous sections, connected through reciprocal bidirectional connections. As in this new model composed of two interacting modules, a full characterization of the bifurcations would require exceedingly high computational cost, we here focused on a representative scenario characterized by two extreme adaptation levels (Fig. 4A). Specifically, the low adaptation level parallels the case of wakefulness as seen in the previous section (case 1, Fig. 1B), as opposed to the high adaptation level characterizing N3 sleep or deep anesthesia (case 4, Fig. 1B). Under conditions of low adaptation, cortical perturbations elicited reverberant interactions between the two modules as reflected by multiple, recurrent and coherent oscillations of evoked activity. Conversely, under conditions of high adaptation, the interplay between the two simulated groups of cortical neurons was short-lasting resulting in only a few oscillations followed by a prominent Off-period, as marked by the suppression of high-frequency power, similar to that observed in sleeping and brain-injured humans. In real-life experiments, the impact of the Off-period on the ability of cortical circuits to sustain causal interactions is assessed by quantifying the duration of the deterministic effects of the perturbation in the phase domain. A key empirical finding is that Off-periods not only disrupt deterministic interaction due to the associated suppression of power but that they also scramble the subsequent phase of the signal when neurons resume firing. Such stochastic reboot is characterized by oscillations with high power but low phase-locking or inter-trial coherence, following the Off-period. We thus performed the same analysis in the model by computing the inter-trial coherence (ITC) averaged across frequencies between 5 and 30 Hz. As shown in Fig. 4A, the Off-period occurring under conditions of high adaptation was followed by a broad-band resumption of power which was however non-deterministic as indicated by a concomitant absence of significant ITC. The main sources of such stochasticity are i) the variable level of fatigue accumulated by the network at the stimulation time and ii) the activity fluctuations due to the finite-size of the neuronal population determining the escape time from the metastable inactive state (Off-period). Notably, in the case of low adaptation, significant deterministic interactions were not curtailed by the Off-period and were longer-lasting. As illustrated in Fig. 4B, these model predictions resulting from the manipulation of the adaptation levels match the power and phase modulations observed in experiments in which TMS or intracranial electrical stimulations are applied in different global brain states, such as during wakefulness, deep NREM sleep and in unresponsive wakefulness syndrome patients (VS/UWS).

**Figure 4.**
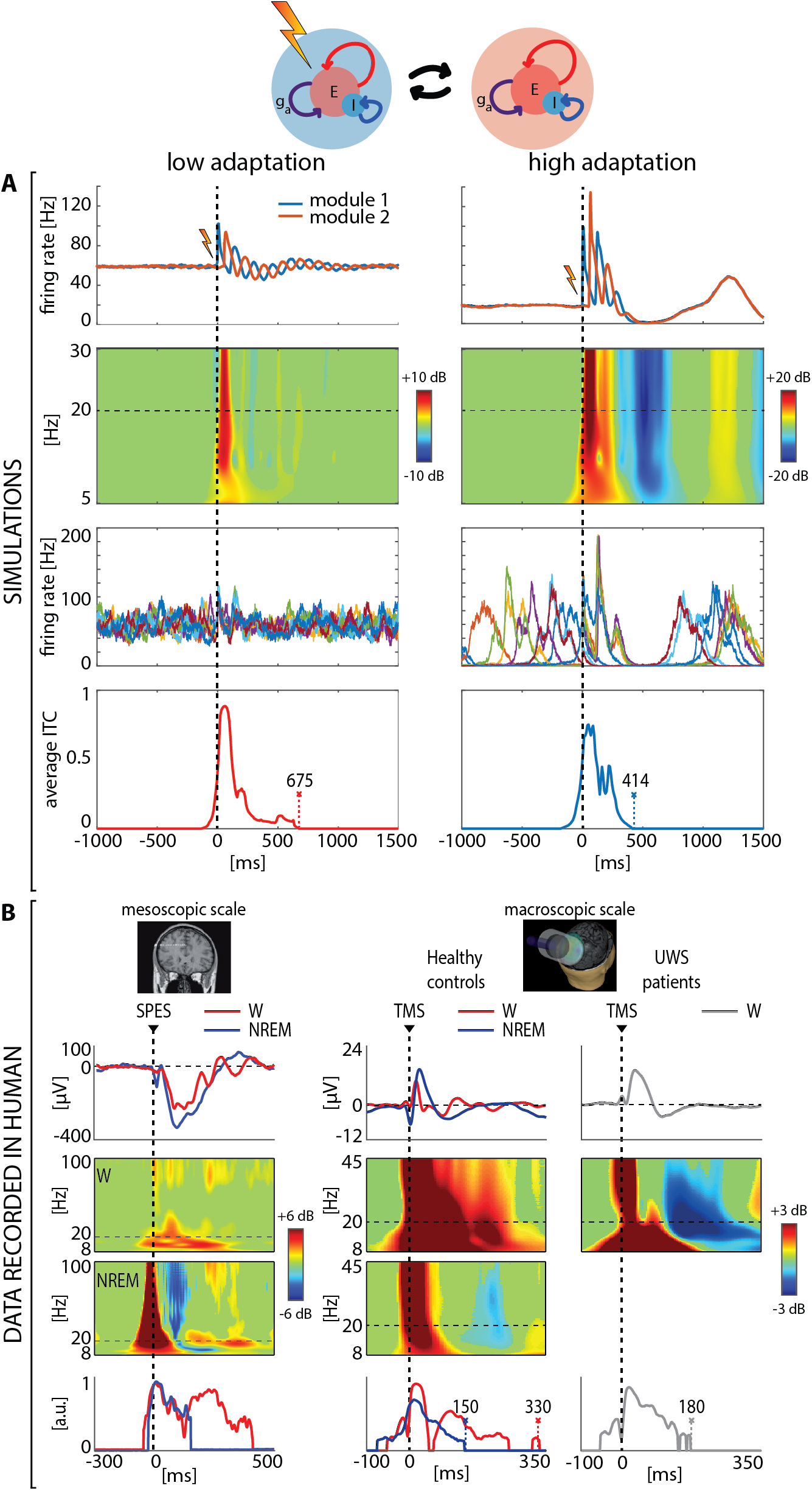
High adaptation level results in early breakdown of causal interactions compared to low adaptation level. **A:** simulated data of two coupled modules for low and high adaptation levels. First row: firing rate activity averaged over 250 trials. A dashed vertical line (at t=0) marks the occurrence of the external stimulus injected into module 1. Second row: Event-related spectral perturbation (ERSP) for module 2. Significance for bootstrap statistics is set at alpha<0.005: absence of significant activation is colored in green, significant increases of power compared to baseline are represented in red, while significant power decreases are colored in blue. Third row: eight firing rate traces of module 2. Fourth row: averaged inter-trial coherence (ITC) for frequencies between 5 and 30 Hz. **B**: Data recorded in human. The first column shows results related to stereo-EEG with SPES during wakefulness (W-red) and NREM sleep (NREM-blue) re-edited from (Pigorini et al., 2015). From top to bottom, average responses of a representative contact, event-related spectral perturbation (ERSP), and phase-locking factor (PLF) for frequencies higher than 8 Hz (details can be found in (Pigorini et al., 2015)). Second and third columns show results related to TMS-EEG re-edited from (Rosanova et al., 2018) encompassing healthy wakefulness (W-red), healthy NREM sleep (NREM-blue) and UWS patients during wakefulness (grey W). As for stereo-EEG-SPES data, from top to bottom, average responses of the channel under the stimulator, event-related spectral perturbation (ERSP), and phase-locking factor (PLF) for frequencies higher than 8 Hz (details can be found in (Rosanova et al., 2018)).

## Discussion

By exploiting a spiking network model endowed with activity-dependent adaptation, the present work provides a principled mechanistic explanation for empirical observations about cortical responsiveness in sleeping healthy subjects, and awake patients with multifocal and focal brain injuries. Specifically, (i) we link changes in cortical reactivity to the presence of underlying activity-dependent adaptation mechanisms, (ii) we highlight the role of perturbations in revealing adaptation mechanisms, and (iii) we expound on the impact of adaptation in disrupting causal interactions among cortical modules.

The first implication of this work is that systematically varying the relationships between excitation and adaptation levels within a formal bifurcation analysis reproduces basic patterns of cortical reactivity to direct perturbations observed in humans. As shown in Fig. 1A and B, increasing levels of adaptation and/or decreasing the excitation engenders a condition whereby cortical circuits react to a direct perturbation with an initial activation followed by an Off-period, yielding responses similar to those found during sleep and in brain-injured patients. This finding offers important elements to interpret macroscale empirical data in terms of neuronal mechanisms.

In this framework, the Off-periods observed after cortical stimulation in sleeping humans can be ascribed to adaptation mechanisms due to slow negative (i.e., inhibitory) feedback produced by calcium- and sodium-gated potassium currents (Sanchez-Vives and McCormick, 2000; Sanchez-Vives et al., 2010), which are enhanced by decreased levels of neuromodulation from brainstem activating systems (McCormick, 1992). Notably, the present simulation results match closely the changes in reactivity to electrical stimulation observed in cortical slices when modulating potassium currents by the application of carbachol and norepinephrine (D’Andola et al., 2018). Recent studies have also pointed to a putative role of active inhibition in conditioning the onset and duration of the Off-period (Funk et al., 2017; Zucca et al., 2017). As adaptation in the present model encompasses self-inhibition, the role of local inhibitory neurons in shaping the changes in cortical responses observed during sleep is not incompatible with the present results.

Interpreting the empirical results obtained in brain-injured and stroke patients within the theoretical framework of the bifurcation diagram discloses a more complex and interesting landscape. Indeed, brain lesions can cause global or local alterations of adaptation and excitation through different mechanisms, even in the context of preserved arousal, as assessed by eyes opening. For example, in some vegetative state patients, lesions, compressions or displacements of brainstem activating systems as well as a critical load of damage to ascending fibers in subcortical white matter may enhance potassium currents (Steriade et al., 1993; Edlow et al., 2012), which corresponds to an upward shift in the bifurcation diagram. In other cases, cortical injuries and white matter lesions can engender a state of cortico-cortical disfacilitation by affecting the excitation term (Takahashi et al., 1981; Lemieux et al., 2014; Rocchi et al., 2022). In this case, the resulting loss of lateral and long-range excitatory input would act by producing shifts on the horizontal axis toward the left, to a point where the probability of evoking an Off-period in an awake patient becomes higher. An interesting implication of the diagram, especially considering the slope of the lower branch of the bifurcation (Fig. 1A and Fig. 3), is that changes in adaptation are expected to have more dramatic effects on cortical bistability as compared to changes in the excitation level. This prediction finds empirical confirmation in cortical slices, an extreme model of cortical injury, whereby reducing adaptation by application of carbachol and norepinephrine is more effective in recovering wake-like responses than increasing excitation by application of kainite (D’Andola et al., 2018). In real-life conditions, however, the two mechanisms (adaptation and excitation) are not mutually exclusive, and may have different relative weights depending on the type/combination of injuries but would both concur in bringing residual cortical circuits into a state in which cortical sleep-like OFF-periods are generated during wakefulness, as observed in stroke (Sarasso et al., 2020) and vegetative state patients (Rosanova et al., 2018).

The second implication of this work lies in the dissociation between observational and perturbational approaches in their ability to unveil the presence of cortical bistability and Off-periods, and bears relevance for the assessment of the aftermath of brain injury. Such dissociation is represented by the area in the excitation-adaptation bifurcation diagram identified in Fig. 3, in which the responses to perturbations more reliably show Off-periods, as compared to spontaneous activity. This portion of the diagram encompasses a relatively small space of the overall dynamics (ranging from wake-like activity to patterns resembling burst suppression) but is extremely relevant in real-life conditions. Indeed, cortical perturbations, either electrical or magnetic, are often capable of evoking clear-cut Off-periods, which are not otherwise present in spontaneous activity not only during N2 sleep but also in many stroke and traumatic brain-injury patients (Rosanova et al., 2018; Sarasso et al., 2020). This is a critical working point along with the sleep-wake transition, in which cortical dynamics are intrinsically unstable as two global activity modes compete (Parrino et al., 2012; Tort-Colet et al., 2021). From a therapeutic perspective, this offers a window of opportunity as small physical or pharmacological perturbation of the network parameters can induce dramatic changes in the brain’s global behavior (Deco et al., 2019). Thus, relating empirical findings to this region of the bifurcation diagram is important for two reasons. First, it suggests that a significant number of patients may lay in a state where the input-output properties of cortical circuits are altered due to a critical, albeit potentially reversible, shift in adaptation (and/or excitation). Second, it shows that, due to its inherent activity-dependent nature, this state of affairs can be better revealed, above and beyond the observation of spontaneous dynamics, by challenging cortical circuits with direct stimulation.

The third result of this theoretical and computational work is that altered levels of adaptation can have profound effects on the capability of cortical neurons to engage in reciprocal interactions as indicated by the complex set of effects observed when changing adaptation in two reciprocally connected cortical modules without modulating their connectivity strength. Under conditions of low adaptation, the two simulated modules engage in a long-lasting series of feed-forward and feed-back interactions leading to multiple waves of activity time-locked to the stimulus, resembling the general pattern of responsiveness found in cortical slices under carbachol and norepinephrine (D’Andola et al., 2018) and in healthy awake subjects (Massimini et al., 2005). For high adaptation, this deterministic pattern of interaction is drastically curtailed; not only Off-periods temporarily obliterate activity but they also disrupt phase-locking to the stimulus once activity resumes. This peculiar condition, whereby high power is associated with minimal levels of phase-locking to the stimulus, is strikingly similar to the pattern found in multiscale empirical measurements ranging from cortical slices (D’Andola et al., 2018) and anesthetized rodents (Arena et al., 2021) to human intracranial and extracranial measurements during NREM sleep and after severe brain injury (Pigorini et al., 2015; Rosanova et al., 2018). Notably, in the brain of patients, Off-periods and the ensuing disruption of causal interactions are empirically associated with low values of whole-brain complexity and with loss of consciousness (Rosanova et al., 2018). Perhaps more importantly, the progressive disappearance of evoked Off-periods is associated with recovery from disorders of consciousness (Rosanova et al., 2018) and stroke (Tscherpel et al., 2020). In light of the present theoretical framework this clinical evolution would correspond to the descending trajectory represented in the bifurcation diagram of Fig. 1A, with potential implications for the stratification, follow-up and rehabilitation in the aftermath of brain injury. For example, detecting cortical bistability by perturbations in a stroke patient points to the presence of functional disruption, adding to the structural damage, and suggests that neuromodulation or pharmacological treatment should aim at reducing adaptation mechanisms and/or strengthening local excitation until the occurrence of evoked Off-period is minimized.

Clearly, the present theoretical framework only represents a first stepping stone upon which more realistic and complex models can be built such as those incorporating the topological organization of cortical networks (Capone et al., 2019; Barbero-Castillo et al., 2021; Pazienti et al., 2022). To be comprehensive such models should also include other subcortical structures like the thalamus whose input is known to play an important role in shaping Off-periods (van Wijngaarden et al., 2016; Zucca et al., 2019). Besides, the present work only provides a minimal account, limited to the proof of principle of two connected modules of the effects of adaptation on cortico-cortical interactions. Hence, a fundamental development will consist in embedding the present framework within a large-scale, connectome-based simulation, such as the “The Virtual Brain”, encompassing a multitude of interacting modules (Sanz Leon et al., 2013; Kringelbach et al., 2020; Goldman et al., 2021; Schirner et al., 2022). This would offer a tool to better understand the effects of local intrusions of cortical bistability within the awake brain, such as those occurring after sleep deprivation (local sleep) (Hung et al., 2013; Sarasso et al., 2014; Bernardi et al., 2015; Nir et al., 2017) as well as those occurring when adaptation and excitation are altered in a regional-specific manner by focal and multifocal structural lesions.

## Acknowledgments

The authors thank Antonio Pazienti, Mario Rosanova, Simone Russo, and Simone Sarasso for engaging with us in crucial discussions during the development of this work, as well as for their comments on the manuscript draft.

## References

Amit DJ, Brunel N (1997) Model of global spontaneous activity and local structured activity during delay periods in the cerebral cortex. Cereb Cortex 7:237–252.

Amit DJ, Tsodyks MV (1991) Quantitative study of attractor neural network retrieving at low spike rates: I. substrate—spikes, rates and neuronal gain. Network: Computation in Neural Systems 2:259–273.

Arena A, Comolatti R, Thon S, Casali AG, Storm JF (2021) General anesthesia disrupts complex cortical dynamics in response to intracranial electrical stimulation in rats. eNeuro:ENEURO.0343-20.2021.

Barbero-Castillo A, Mateos-Aparicio P, Dalla Porta L, Camassa A, Perez-Mendez L, Sanchez-Vives MV (2021) Impact of GABAA and GABAB Inhibition on Cortical Dynamics and Perturbational Complexity during Synchronous and Desynchronized States. J Neurosci 41:5029–5044.

Bernardi G, Siclari F, Yu X, Zennig C, Bellesi M, Ricciardi E, Cirelli C, Ghilardi MF, Pietrini P, Tononi G (2015) Neural and behavioral correlates of extended training during sleep deprivation in humans: evidence for local, task-specific effects. J Neurosci 35:4487– 4500.

Brunel N, Hakim V (1999) Fast global oscillations in networks of integrate-and-fire neurons with low firing rates. Neural Comput 11:1621–1671.

Capocelli RM, Ricciardi LM (1971) Diffusion approximation and first passage time problem for a model neuron. Kybernetik 8:214–223.

Capone C, Rebollo B, Muñoz A, Illa X, Del Giudice P, Sanchez-Vives MV, Mattia M (2019) Slow Waves in Cortical Slices: How Spontaneous Activity is Shaped by Laminar Structure. Cereb Cortex 29:319–335.

Cash SS, Halgren E, Dehghani N, Rossetti AO, Thesen T, Wang C, Devinsky O, Kuzniecky R, Doyle W, Madsen JR, Bromfield E, Eross L, Halász P, Karmos G, Csercsa R, Wittner L, Ulbert I (2009) The human K-complex represents an isolated cortical down-state. Science 324:1084–1087.

Chang J-Y, Pigorini A, Massimini M, Tononi G, Nobili L, Van Veen BD (2012) Multivariate autoregressive models with exogenous inputs for intracerebral responses to direct electrical stimulation of the human brain. Front Hum Neurosci 6:317.

Compte A, Sanchez-Vives MV, McCormick DA, Wang X-J (2003) Cellular and network mechanisms of slow oscillatory activity (<1 Hz) and wave propagations in a cortical network model. J Neurophysiol 89:2707–2725.

D’Andola M, Rebollo B, Casali AG, Weinert JF, Pigorini A, Villa R, Massimini M, Sanchez-Vives MV (2018) Bistability, Causality, and Complexity in Cortical Networks: An In Vitro Perturbational Study. Cereb Cortex 28:2233–2242.

Deco G, Cruzat J, Cabral J, Tagliazucchi E, Laufs H, Logothetis NK, Kringelbach ML (2019) Awakening: Predicting external stimulation to force transitions between different brain states. Proc Natl Acad Sci U S A 116:18088–18097.

Delorme A, Makeig S (2004) EEGLAB: an open source toolbox for analysis of single-trial EEG dynamics including independent component analysis. J Neurosci Methods 134:9–21.

Dhooge A, Govaerts W, Kuznetsov YuA (2003) MATCONT: A MATLAB package for numerical bifurcation analysis of ODEs. ACM Trans Math Softw 29:141–164.

Edlow BL, Takahashi E, Wu O, Benner T, Dai G, Bu L, Grant PE, Greer DM, Greenberg SM, Kinney HC, Folkerth RD (2012) Neuroanatomic connectivity of the human ascending arousal system critical to consciousness and its disorders. J Neuropathol Exp Neurol 71:531–546.

Funk CM, Peelman K, Bellesi M, Marshall W, Cirelli C, Tononi G (2017) Role of Somatostatin-Positive Cortical Interneurons in the Generation of Sleep Slow Waves. J Neurosci 37:9132–9148.

Gerstner W, Kistler WM, Naud R, Paninski L (2014) Neuronal Dynamics: From Single Neurons To Networks And Models Of Cognition. undefined Available at: https://www.semanticscholar.org/paper/Neuronal-Dynamics%3A-From-Single-Neurons-To-Networks-Gerstner-Kistler/c9e81db0895b027a0248acf38d5149e0c492b070 [Accessed May 11, 2022].

Gigante G, Mattia M, Giudice PD (2007) Diverse Population-Bursting Modes of Adapting Spiking Neurons. Phys Rev Lett 98:148101.

Goldman JS, Kusch L, Yalçinkaya BH, Depannemaecker D, Nghiem T-AE, Jirsa V, Destexhe A (2021) A comprehensive neural simulation of slow-wave sleep and highly responsive wakefulness dynamics. Neuroscience. Available at: http://biorxiv.org/lookup/doi/10.1101/2021.08.31.458365 [Accessed April 8, 2022].

Hung C-S, Sarasso S, Ferrarelli F, Riedner B, Ghilardi MF, Cirelli C, Tononi G (2013) Local experience-dependent changes in the wake EEG after prolonged wakefulness. Sleep 36:59–72.

Kringelbach ML, Cruzat J, Cabral J, Knudsen GM, Carhart-Harris R, Whybrow PC, Logothetis NK, Deco G (2020) Dynamic coupling of whole-brain neuronal and neurotransmitter systems. Proc Natl Acad Sci U S A 117:9566–9576.

Kuznetsov Y (2004) Elements of Applied Bifurcation Theory, 3rd ed. New York: Springer-Verlag. Available at: https://www.springer.com/gp/book/9780387219066 [Accessed June 30, 2021].

Latham PE, Richmond BJ, Nelson PG, Nirenberg S (2000) Intrinsic dynamics in neuronal networks. I. Theory. J Neurophysiol 83:808–827.

Lemieux M, Chen J-Y, Lonjers P, Bazhenov M, Timofeev I (2014) The impact of cortical deafferentation on the neocortical slow oscillation. J Neurosci 34:5689–5703.

Lewis LD, Weiner VS, Mukamel EA, Donoghue JA, Eskandar EN, Madsen JR, Anderson WS, Hochberg LR, Cash SS, Brown EN, Purdon PL (2012) Rapid fragmentation of neuronal networks at the onset of propofol-induced unconsciousness. Proc Natl Acad Sci U S A 109:E3377–3386.

Mascaro M, Amit DJ (1999) Effective neural response function for collective population states. Network 10:351–373.

Massimini M, Ferrarelli F, Esser SK, Riedner BA, Huber R, Murphy M, Peterson MJ, Tononi G (2007) Triggering sleep slow waves by transcranial magnetic stimulation. Proc Natl Acad Sci U S A 104:8496–8501.

Massimini M, Ferrarelli F, Huber R, Esser SK, Singh H, Tononi G (2005) Breakdown of cortical effective connectivity during sleep. Science 309:2228–2232.

Mattia M, Biggio M, Galluzzi A, Storace M (2019) Dimensional reduction in networks of non-Markovian spiking neurons: Equivalence of synaptic filtering and heterogeneous propagation delays. PLoS Comput Biol 15:e1007404.

Mattia M, Del Giudice P (2002) Population dynamics of interacting spiking neurons. Phys Rev E 66:051917.

Mattia M, Giudice PD (2000) Efficient Event-Driven Simulation of Large Networks of Spiking Neurons and Dynamical Synapses. Neural Computation 12:2305–2329.

Mattia M, Sanchez-Vives MV (2012) Exploring the spectrum of dynamical regimes and timescales in spontaneous cortical activity. Cogn Neurodyn 6:239–250.

McCormick DA (1992) Neurotransmitter actions in the thalamus and cerebral cortex and their role in neuromodulation of thalamocortical activity. Prog Neurobiol 39:337– 388.

Mukovski M, Chauvette S, Timofeev I, Volgushev M (2007) Detection of active and silent states in neocortical neurons from the field potential signal during slow-wave sleep. Cereb Cortex 17:400–414.

Nir Y, Andrillon T, Marmelshtein A, Suthana N, Cirelli C, Tononi G, Fried I (2017) Selective neuronal lapses precede human cognitive lapses following sleep deprivation. Nat Med 23:1474–1480.

Nir Y, Staba RJ, Andrillon T, Vyazovskiy VV, Cirelli C, Fried I, Tononi G (2011) Regional slow waves and spindles in human sleep. Neuron 70:153–169.

Parrino L, Ferri R, Bruni O, Terzano MG (2012) Cyclic alternating pattern (CAP): the marker of sleep instability. Sleep Med Rev 16:27–45.

Pazienti A, Galluzzi A, Dasilva M, Sanchez-Vives MV, Mattia M (2022) Slow waves form expanding, memory-rich mesostates steered by local excitability in fading anesthesia. iScience 25:103918.

Piantoni G, Astill RG, Raymann RJEM, Vis JC, Coppens JE, Van Someren EJW (2013) Modulation of γ and spindle-range power by slow oscillations in scalp sleep EEG of children. Int J Psychophysiol 89:252–258.

Pigorini A, Sarasso S, Proserpio P, Szymanski C, Arnulfo G, Casarotto S, Fecchio M, Rosanova M, Mariotti M, Lo Russo G, Palva JM, Nobili L, Massimini M (2015) Bistability breaks-off deterministic responses to intracortical stimulation during non-REM sleep. Neuroimage 112:105–113.

Richardson MJE, Swarbrick R (2010) Firing-rate response of a neuron receiving excitatory and inhibitory synaptic shot noise. Phys Rev Lett 105:178102.

Rocchi F, Canella C, Noei S, Gutierrez-Barragan D, Coletta L, Galbusera A, Stuefer A, Vassanelli S, Pasqualetti M, Iurilli G, Panzeri S, Gozzi A (2022) Increased fMRI connectivity upon chemogenetic inhibition of the mouse prefrontal cortex. Nat Commun 13:1056.

Rosanova M, Fecchio M, Casarotto S, Sarasso S, Casali AG, Pigorini A, Comanducci A, Seregni F, Devalle G, Citerio G, Bodart O, Boly M, Gosseries O, Laureys S, Massimini M (2018) Sleep-like cortical OFF-periods disrupt causality and complexity in the brain of unresponsive wakefulness syndrome patients. Nat Commun 9:4427.

Sanchez-Vives MV, Mattia M, Compte A, Perez-Zabalza M, Winograd M, Descalzo VF, Reig R (2010) Inhibitory modulation of cortical up states. J Neurophysiol 104:1314–1324.

Sanchez-Vives MV, McCormick DA (2000) Cellular and network mechanisms of rhythmic recurrent activity in neocortex. Nat Neurosci 3:1027–1034.

Sanz Leon P, Knock SA, Woodman MM, Domide L, Mersmann J, McIntosh AR, Jirsa V (2013) The Virtual Brain: a simulator of primate brain network dynamics. Front Neuroinform 7:10.

Sarasso S, D’Ambrosio S, Fecchio M, Casarotto S, Viganò A, Landi C, Mattavelli G, Gosseries O, Quarenghi M, Laureys S, Devalle G, Rosanova M, Massimini M (2020) Local sleep-like cortical reactivity in the awake brain after focal injury. Brain 143:3672–3684.

Sarasso S, Pigorini A, Proserpio P, Gibbs SA, Massimini M, Nobili L (2014) Fluid boundaries between wake and sleep: experimental evidence from Stereo-EEG recordings. Arch Ital Biol 152:169–177.

Schirner M et al. (2022) Brain simulation as a cloud service: The Virtual Brain on EBRAINS. Neuroimage 251:118973.

Siegert AJF (1951) On the First Passage Time Probability Problem. Phys Rev 81:617–623.

Steriade M, Nuñez A, Amzica F (1993) A novel slow (< 1 Hz) oscillation of neocortical neurons in vivo: depolarizing and hyperpolarizing components. J Neurosci 13:3252– 3265.

Takahashi H, Manaka S, Sano K (1981) Changes in extracellular potassium concentration in cortex and brain stem during the acute phase of experimental closed head injury. J Neurosurg 55:708–717.

Tononi G, Massimini M (2008) Why does consciousness fade in early sleep? Ann N Y Acad Sci 1129:330–334.

Torao-Angosto M, Manasanch A, Mattia M, Sanchez-Vives MV (2021) Up and Down States During Slow Oscillations in Slow-Wave Sleep and Different Levels of Anesthesia. Front Syst Neurosci 15:609645.

Tort-Colet N, Capone C, Sanchez-Vives MV, Mattia M (2021) Attractor competition enriches cortical dynamics during awakening from anesthesia. Cell Rep 35:109270.

Treves A (1993) Mean-field analysis of neuronal spike dynamics. Network:259–284.

Tscherpel C, Dern S, Hensel L, Ziemann U, Fink GR, Grefkes C (2020) Brain responsivity provides an individual readout for motor recovery after stroke. Brain 143:1873– 1888.

Tuckwell HC (1988) Introduction to Theoretical Neurobiology: Volume 2: Nonlinear and Stochastic Theories. Cambridge: Cambridge University Press. Available at: https://www.cambridge.org/core/books/introduction-to-theoretical-neurobiology/5B00C4410746818CE64451F201CD1FE7 [Accessed May 16, 2022].

Usami K, Matsumoto R, Kobayashi K, Hitomi T, Shimotake A, Kikuchi T, Matsuhashi M, Kunieda T, Mikuni N, Miyamoto S, Fukuyama H, Takahashi R, Ikeda A (2015) Sleep modulates cortical connectivity and excitability in humans: Direct evidence from neural activity induced by single-pulse electrical stimulation. Hum Brain Mapp 36:4714–4729.

van Wijngaarden JBG, Zucca R, Finnigan S, Verschure PFMJ (2016) The Impact of Cortical Lesions on Thalamo-Cortical Network Dynamics after Acute Ischaemic Stroke: A Combined Experimental and Theoretical Study. PLoS Comput Biol 12:e1005048.

Zucca S, D’Urso G, Pasquale V, Vecchia D, Pica G, Bovetti S, Moretti C, Varani S, Molano-Mazón M, Chiappalone M, Panzeri S, Fellin T (2017) An inhibitory gate for state transition in cortex. Elife 6:e26177.

Zucca S, Pasquale V, Lagomarsino de Leon Roig P, Panzeri S, Fellin T (2019) Thalamic Drive of Cortical Parvalbumin-Positive Interneurons during Down States in Anesthetized Mice. Curr Biol 29:1481-1490.e6.

