## Supplementary material for "Adaptation shapes local cortical reactivity: from bifurcation diagram and simulations to human physiological and pathological responses": Figure 1-1

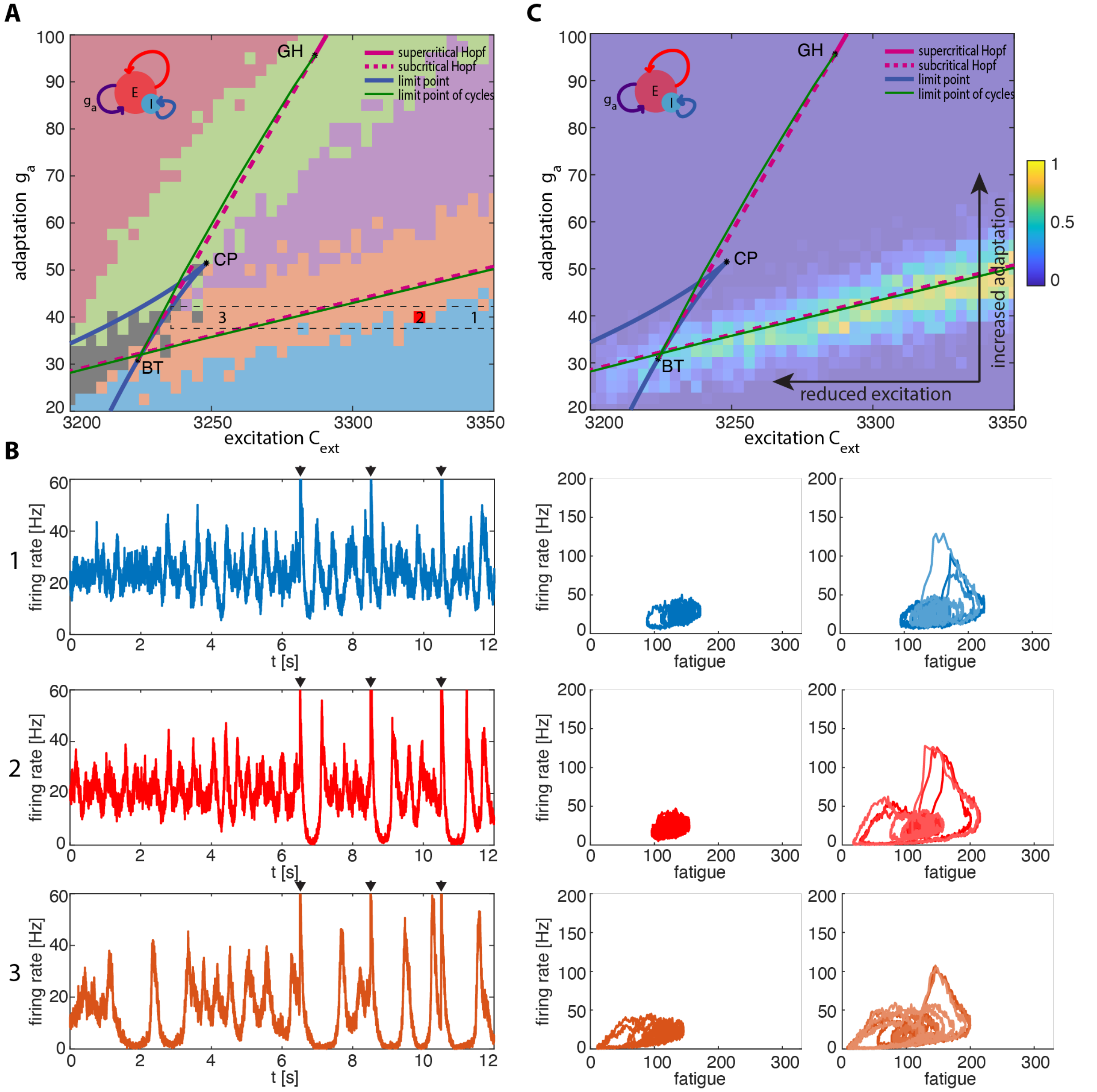


**Figure 1-1**. Bifurcation analysis of the population rate model, and dynamical regimes of the spiking neuron network (as in Fig. 1) for a fixed adaptation level (g = 42.5) and different excitation levels (3342.5, 3320, 3248.75). **A** and **C** as in Fig. 1 and Fig. 3, respectively. **B:** Spontaneous and stimulus-evoked signals (using the same color-coding as in A) for a fixed level of adaptation. Left column, time series encompassing both spontaneous activity (up to 6000 time steps) and stimulus-evoked activity (remaining interval). Black triangles indicate the occurrence of the stimulation. Central column, spontaneous activity as a function of fatigue. Right column, three superimposed orbits due to perturbations as a function of the fatigue.
